# Default But Not Rest: Topological Discrimination Defines the Default Mode Network

**DOI:** 10.1101/2022.11.10.516071

**Authors:** Bo Wang, Tiangang Zhou, Sheng He, Lin Chen

## Abstract

The default mode network (DMN), a set of transmodal cortical regions, has historically been argued to serve the internal functions of brain. The discovery of this network highlighted the brain’s intrinsic operations. The DMN generally decreases its activity during tasks and increases its activity during relaxed non-task states. It is important to investigate the nature of the DMN in order to understand the human brain in health and disease. In the current study, we discovered a task-related cortical network we called the Topological Discrimination Network (TDN), which was consistently revealed by contrasting activations from topological discrimination tasks with local geometric discrimination tasks. The TDN and the DMN consist of essentially the same group of brain regions and the fMRI response of topological discrimination in those regions exhibited consistent temporal dynamics with resting state. The robustness of the TDN is supported by multiple experiments performed at different field strengths (3T and 7T MRI scanner) as well as different types of signals measured (BOLD and CBF). The collective results suggest that the process of topological discrimination could almost be considered as a functional “default mode” of our brain. The TDN, like the DMN, could define the functional baseline of brain, with the advantage of functional consistency across participants and experimental sessions.

## Introduction

The default mode network (DMN) has been identified reliably during rest in the human, nonhuman primate, cat, and even rodent brains. It is typically considered a task-negative network due to the reduction of its activity across a broad-spectrum of tasks (Gordon L. Shulman, 1997; Raichle et al., 2001). The discovery of the DMN provides a more balanced view of the brain, highlighting the importance of the brain’s organized intrinsic ongoing activity, in addition to its reflexive functions (Raichle, 2006, 2010, 2011; Raichle, 2015a, 2015b; Raichle et al., 2001). Although the DMN was characterized as “task negative” in the earlier studies, it also supports some mental operations such as self-reference (Davey, Pujol, & Harrison, 2016; D. A. Gusnard, Akbudak, Shulman, & Raichle, 2001), working memory (Sormaz et al., 2018; Vatansever, Menon, Manktelow, Sahakian, & Stamatakis, 2015), mental time traveling (Ylva Østby, 2012), mind-wandering (Andrews-Hanna, 2012; Buckner, Andrews-Hanna, & Schacter, 2008; Sormaz et al., 2018). However, the commonalities between these functions are still unclear (Murphy et al., 2018) and the functions of the DMN remain a puzzle (Raichle, 2015a).

The discovery of the DMN made apparent the need for additional ways to study the large-scale intrinsic functional organization of the brain. In the past two decades, investigators had been trying to characterize the spatiotemporal structure of this intrinsic activity by studying the task-unrelated, seemingly spontaneous signals of brain. However, because of the very unconstrained nature of the task-unrelated signals, it was difficult to interpret them. It might be more productive to adopt designs with enhanced control and interpretability while respect the importance of intrinsic activity (Finn, 2021). A promising approach was mapping this functional organization by comparison of cortical activities under some specific cognitive processes which were better controlled.

The topological approach to perceptual organization claims that the perception of topological properties is a primitive and general function of our visual system, proposed as the theory of “global-first” (Chen, 1982, 2005; Chen, Zhang, & Srinivasan, 2003; He, Zhou, Zhou, He, & Chen, 2015; B. Wang, Zhou, Zhuo, & Chen, 2007; Zhou, Luo, Zhou, Zhuo, & Chen, 2010; Zhuo et al., 2003). The time dependence of perceiving form properties is systematically related to their structural stability under change; in particular, the perception of local geometrical properties depends on topological perception (based on physical connectivity), and topological perception occurs earlier than the perception of local geometrical properties (Chen, 2005). Based on these considerations, it is possible that the comparison between topological perception with perception of local geometrical properties could serve as a functional baseline of the brain. The present fMRI study aimed at revealing the brain activations in response to a visual discrimination task of topological properties vs. local geometrical tasks, and comparing the corresponding network with the DMN.

In this fMRI study, we performed 6 fMRI experiments to investigate the different brain responses to topological discrimination and local geometrical perception systematically, in comparison to the resting state activities.

## Topological Discrimination Evoked DMN-like BOLD Responses at 3T

### EXPERIMENT 1

#### METHODS

##### Subjects

10 males and 11 females, between 19 to 28 years old, participated in Experiment 1 as paid volunteers. All of them had normal or corrected-to-normal visual acuity and no history of neurological or psychiatric illness. They all provided informed consent approved by the local ethics committee of Beijing MRI Center for Brain Research (BMCBR).

#### Stimuli and Procedure

The experimental paradigm was generated based on the “configural superiority effects” (Pomerantz, Sager, & Stoever, 1977), which refer to the findings that configural relations between simple components (line segments in figures) rather than the components (line segments) themselves. In the test of configural effects, subjects were always asked to report which quadrant differs (disparate one) from the other three in a four-quadrant stimulus (Chen, 2005).

Four sets of stimuli were adopted in this experiment (Figure 1A). Stimulus ***a*** served as a discrimination task based on orientation of arrows, a kind of Euclidean property. Stimulus ***b*** illustrated topological discrimination based on the difference in holes by using the “triangle-arrow pair” figure. The triangle in the disparate quadrant was made up of exactly the same three line-segments as the arrows in the rest of the quadrants, but its closed nature makes it topologically different from the arrows. Stimulus ***c*** represents texture discrimination based on a difference in orientation of angles, also a kind of Euclidean property. But each quadrant contains exactly the same line segments. Stimulus ***d*** illustrated topological discrimination too. It was designed as “square-angle pair” figure, in which the disparate quadrant contains a larger square formed by four right angles, but the rest each contain four right angles distributed in random orientations.

**Figure 1.**
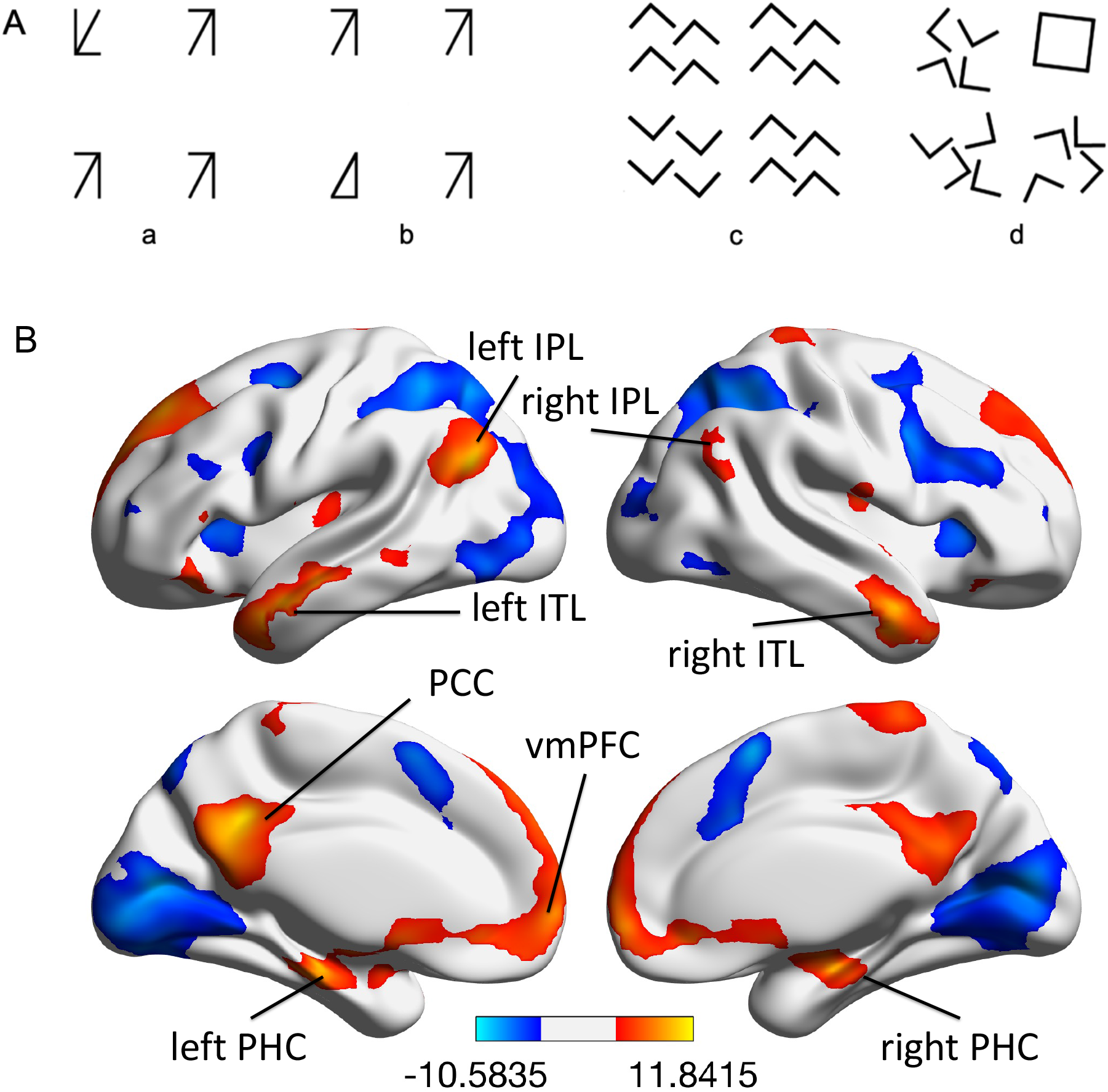
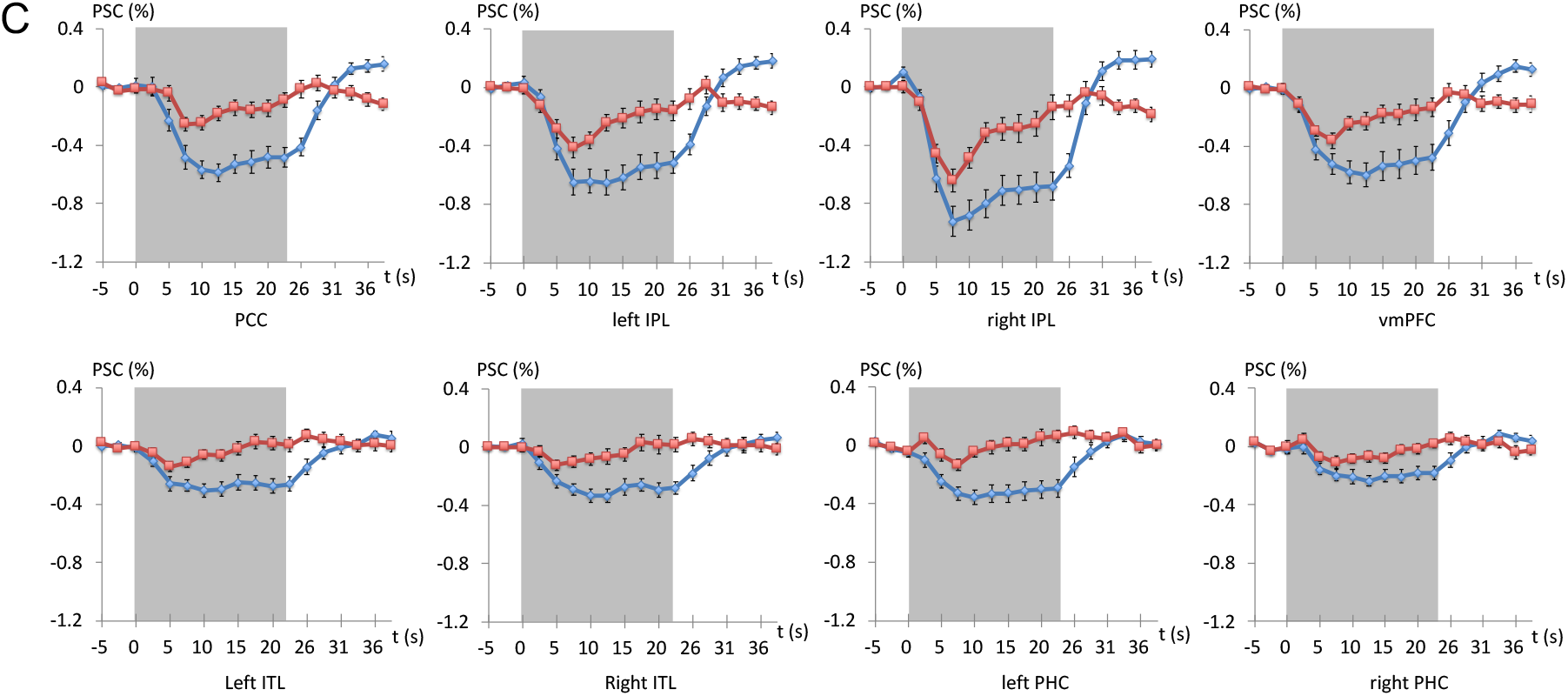
The stimuli and the results of Experiment 1. (A) Four sets of stimuli; (B) comparison of fMRI activations between the topological and local tasks. (C) The averaged BOLD signals responding to topological (red) or local (blue) task in eight major ROIs. The gray box indicated the time window of the task blocks.

Visual stimuli were projected on a rear-projection screen placed 70 cm from the subject’s eyes in the bore of the scanner and viewed through an angled mirror attached to the head coil. All stimuli were in black on a gray background. The small figures in each quadrant and the whole stimulus were displayed in size of 2.0° × 1.2° and 8° × 6°, respectively. The experiment was coded with Psychtoolbox implemented in MATLAB (www.mathworks.com) on a Windows XP computer.

Participants completed a total of three functional runs (6 min 38 secs per run) in the scanner. Within each run, there were two blocks related to each of the 4 sets of stimuli. Each block consisted of 8 trials and lasted for 25.6 seconds. In each trial, the stimulus display was presented at the center of the screen for 1 second and the odd small figure was equally likely (25% chance) to appear in any one of the four quadrants. Participants were instructed, with emphasis, to find the disparate quadrant and report its position while the stimulus display appeared as accurately and quickly as possible. No feedback was given.

#### MR image acquisition and data analysis

Functional MRI was acquired at a Siemens 3T Trio Tim scanner (Erlangen, Germany) equipped with one12-channel head coils. Functional images were acquired using a gradient echo-planar imaging (EPI) sequence [TR = 2,560 ms; TE = 30 ms; FOV= 220 × 220 mm^2^; flip angle = 90°; matrix size = 64 × 64; 48 transversal slices along the AC–PC plane, slice thickness = 2.4 mm^3^, gap = 0.6 mm; parallel imaging with SENSE = 2]. For each task run, a total of 132 EPI volume images were acquired. High-resolution T1-weighted 3D-MPRAGE images were acquired (TR/TE/TI = 2,530 ms/1.64 ms/1,100 ms; flip angle = 7°; 176 sagittal slices; field of view = 256 × 256 mm^2^; resolution = 1.0 × 1.0 × 1.0 mm^3^) for registration of functional images and localization of brain activation.

fMRI data were aligned, unwarping for distortion correction, co-registered to the T1 structure image, spatially normalized to the MNI space, and smoothed by an 8-mm FWHM Gaussian filter using SPM12 (Wellcome Trust Centre for Neuroimaging, London, www.fil.ion.ucl.ac.uk/spm/) after the first two volumes of each functional run were discarded to allow for magnetization equilibration. At the first level of analysis, contrast images of topological vs. local discriminations was calculated on each subject’s data by using the General Linear Model (GLM). These contrasts images then were introduced into the second-level random-effect analysis to allow for population inferences. One-sample t test was conducted to evaluate group activation maps of the topological discrimination responses. The data of the stimuli which represented topological (stimulus ***b & d***) or local (stimulus ***a & c***) geometrical difference were combined respectively to obtained convergent comparison of two discrimination tasks with solider controlling for stimuli, basic sensory, motor and attentional influences.

To quantify the functional response to the different tasks, we computed the averaged time course of the percent signal changes (PSCs) of BOLD signal in each activation regions across all subjects and repetitions.

## RESULTS

Figure 1B shows the activation map of the group level comparison between the topological discrimination and local discrimination task (stimulus ***b & d*** vs. stimulus ***a & c***). The topological discrimination revealed increased activity in multiple cortical regions, including the posterior cingulate cortex (PCC), the ventral medial prefrontal cortex (vmPFC), the bilateral inferior parietal lobule (IPL), the bilateral inferior temporal lobe (ITL, anterolateral part of the temporal cortex), and the bilateral parahippocampal cortex (PHC). Remarkably, all of those regions passed FWE correction and fell in the DMN (Table 1).

**Table 1.**
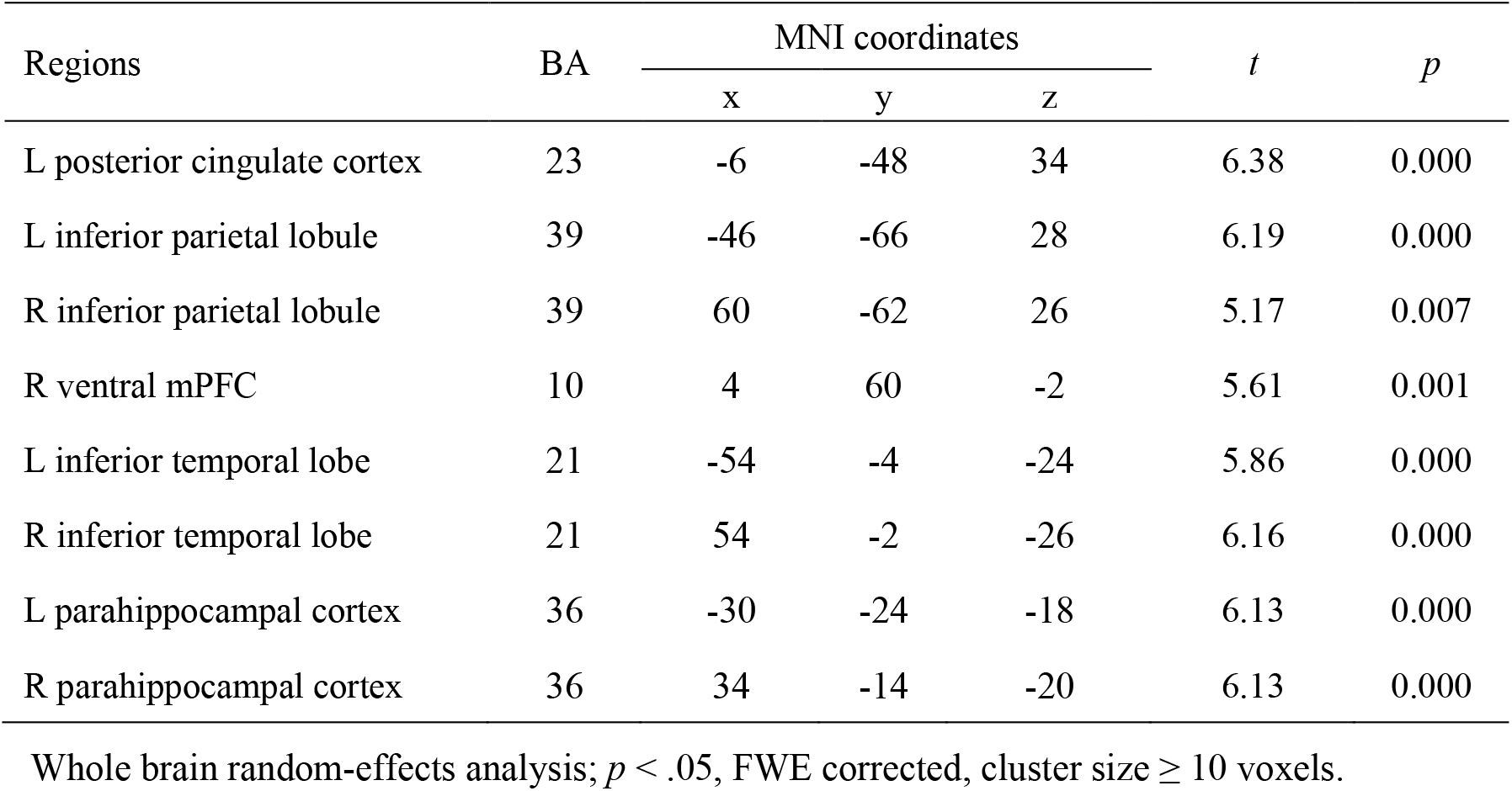
The active regions of the topological vs. local geometrical discrimination

| Regions | BA | MNI coordinates |  |  | *t* | *p* |
| --- | --- | --- | --- | --- | --- | --- |
|  |  | x | y | z |  |  |
| L posterior cingulate cortex | 23 | -6 | -48 | 34 | 6.38 | 0.000 |
| L inferior parietal lobule | 39 | -46 | -66 | 28 | 6.19 | 0.000 |
| R inferior parietal lobule | 39 | 60 | -62 | 26 | 5.17 | 0.007 |
| R ventral mPFC | 10 | 4 | 60 | -2 | 5.61 | 0.001 |
| L inferior temporal lobe | 21 | -54 | -4 | -24 | 5.86 | 0.000 |
| R inferior temporal lobe | 21 | 54 | -2 | -26 | 6.16 | 0.000 |
| L parahippocampal cortex | 36 | -30 | -24 | -18 | 6.13 | 0.000 |
| R parahippocampal cortex | 36 | 34 | -14 | -20 | 6.13 | 0.000 |

Whole brain random-effects analysis; $p < .05$ , FWE corrected, cluster size $\geq 10$ voxels.

As shown in Figure 1C, the time course of the BOLD response to the topological discrimination had no significantly (the bilateral ITL and the bilateral PHC, *p* = .467; *p* = .504; *p* = .910, *p* =0.131) or minor lower difference compared with the fixation (resting) period before task block in all the eight active regions. In contrast, the discrimination of local geometrical stimulus evoked significant task induced deactivation (TID) in all regions in comparison with the resting (*p* < .001).

### EXPERIMENT 2

In Experiment 1, we revealed a DMN-like brain activation pattern, showing greater responses to topological discrimination than local discrimination based on two pairs of stimuli. In Experiment 2, we extended the discrimination task with 6 sets of stimuli to investigate the generalizability of brain responses to topological discrimination tasks.

#### METHODS

##### Subjects

13 males and 19 females, between 18 to 28 years old, participated in Experiment 2 as paid volunteers.

#### Stimuli and Procedure

Six sets of stimuli were systematically measured as one varies form properties at different levels of geometrical properties: In turn, the differences in orientation of angles (a kind of Euclidean property) (Figure 2Aa), parallelism (a kind of affine property) (Figure 2Ab), collinearity (a kind of projective property) (Figure 2Ac), and holes (Figure 2Ad & Ae).

**Figure 2.**
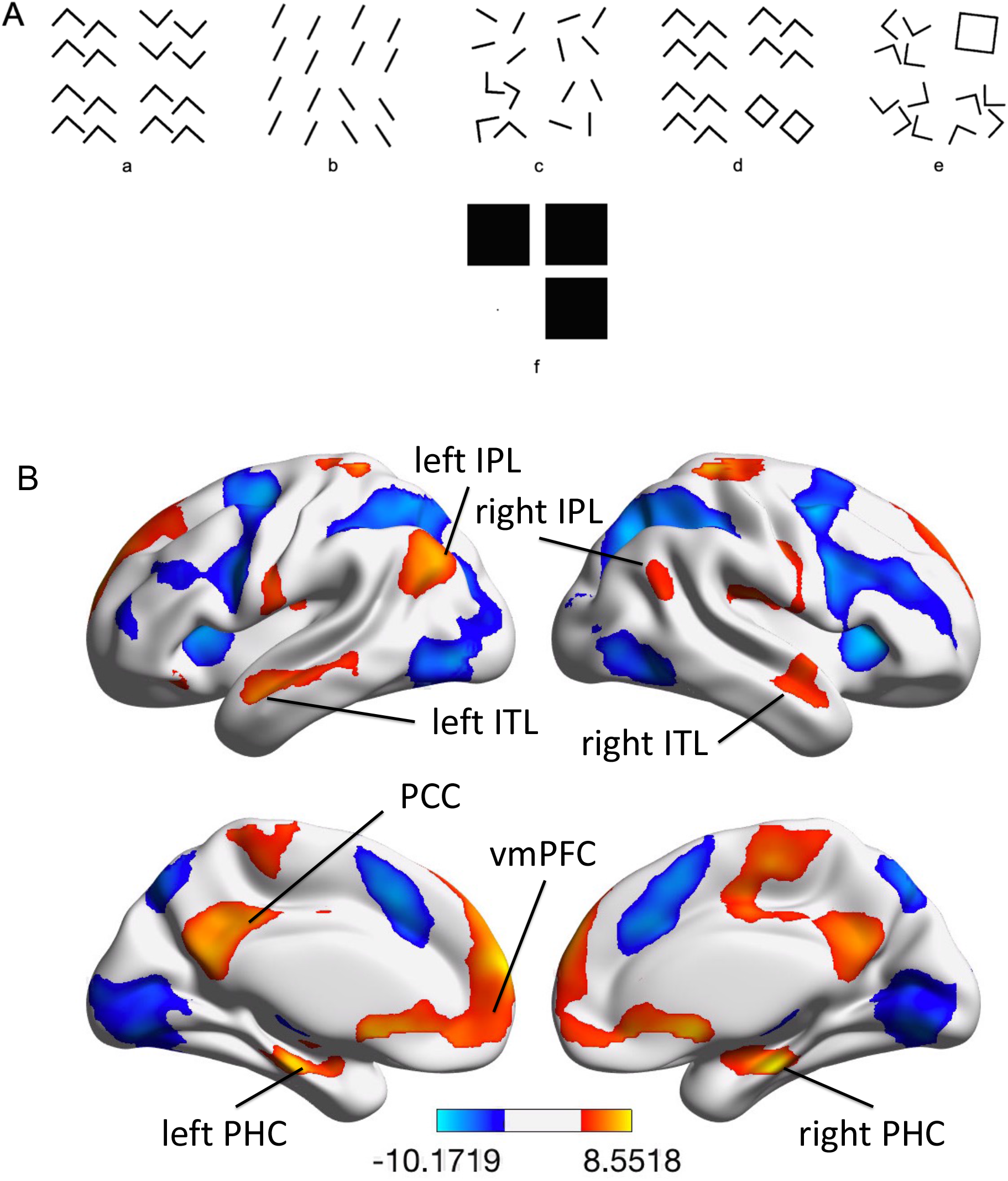
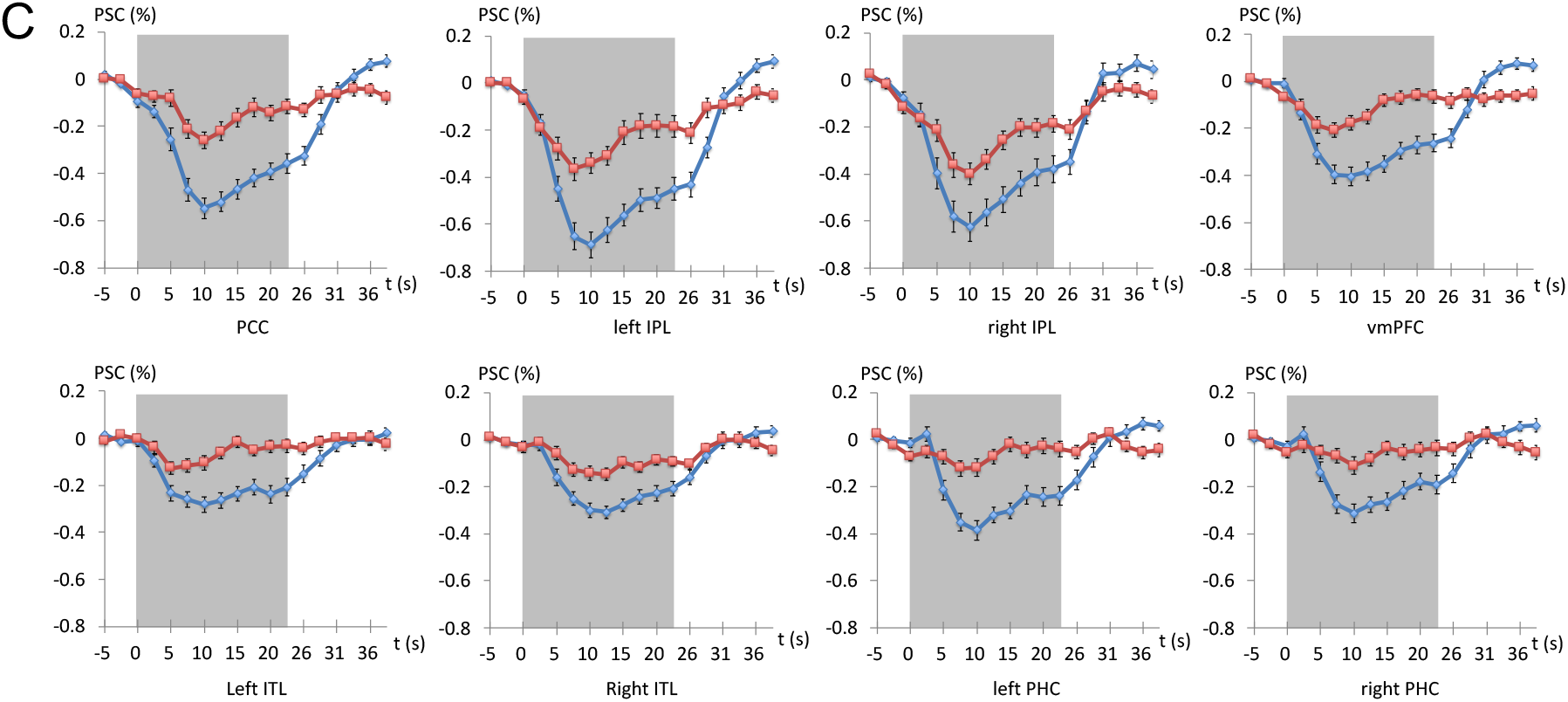
The stimuli and the results of Experiment 2. (A) Six sets of stimuli; (B) comparison of fMRI activations between the topological and local discrimination tasks. (C) The averaged PSCs responding to the topological (red) and local (blue) tasks in the eight main ROIs. The gray box indicated the temporal duration of task blocks.

Stimuli ***a*** and ***e*** were used in Experiment 1. In stimulus ***b***, the line segments in the disparate quadrant are parallel but differ in parallelism from the rest. In stimulus ***c***, the disparate quadrant contains bent lines while there are straight lines in the remaining quadrants. Stimulus ***d*** was adapted from stimulus ***a***: in its disparate quadrant, two of the four right angles are rotated 180° so that they could join another two right angles to form two squares, while the four right angles in each of the rest quadrants are open.

Stimulus ***f*** (Figure 2Af) was designed in such a way that the disparate quadrant contains no figure, whereas the remaining quadrants each contain a large solid square. One can hardly imagine a task easier than this disparate quadrant task; therefore, it should be used to establish a baseline for the easiest task. Moreover, the topological approach leads to a novel analysis of the nature of the baseline: The property of presence vs. absence may also be considered a kind of topological invariant, because topological transformations neither create nor destroy objects. Hence, despite their great difference in local features, Figure 2Ad, Ae and Af share the same intrinsic characteristic of topological difference.

The small figure in each quadrant and the whole stimulus were displayed in size of 3.3° × 3.3° and 10° × 10°, respectively. The background screen, the colors of those stimuli, the experimental equipment and software were the same as in Experiment 1.

Participants completed seven functional runs (5 min 38 secs per run) after an 8-min resting scan. Within each functional run, there were six blocks (one block for each of the 6 sets of stimuli). Each block contained eight 3.2-second trials. The task and requirements for participants were the same as in Experiment 1.

#### MR image acquisition and analysis

The scanner, sequences and all peripheral equipment used in this experiment were all the same with Experiment 1, except that a 20-channel receiver head coil was used instead of the 12-channal one.

The voxel wise analysis and the PSCs calculation procedures were the same as in Experiment 1 for task fMRI data. In addition, a functional connectivity (FC) analysis of the resting state fMRI (rs-fMRI) data was performed by using Data Processing Assistant for Brain Imaging (DPABI) (Yan, Wang, Zuo, & Zang, 2016). The first 10 volumes of images were discarded before slice timing and head motion correction. Then, all the images were normalized to the MNI space, resampled at 2 × 2 × 2 mm^3^ resolution; applied linear trend removal, band-pass filtered (0.01–0.1 Hz) and spatially smoothed (6 mm FWHM Gaussian Kernel). Then, some nuisance covariates were regressed out, including rigid-body 6 head motion parameters, white matter signal, and cerebrospinal fluid signal. The residual time series were used in the subsequent network analysis. Finally, the PCC was defined as the seed region for a seed-based FC analyses to map the DMN. In the statistical map at group level across all subjects, the major hubs of the DMN were defined as regions of interest (ROIs) for the ROI analysis on the task data in the following experiments.

## RESULTS

### Task fMRI

The result of the comparison between the topological and local discrimination tasks (Figure 2A, stimulus ***d, e & f*** vs. stimulus ***a, b & c***) revealed a consistent DMN-like activation patterns as in Experiment 1. All of the active regions passed FWE correction (Table 2).

**Table 2.** The active regions of the topological vs. local discrimination

| Regions | BA | MNI coordinates |  |  | *t* | *p* |
| --- | --- | --- | --- | --- | --- | --- |
|  |  | x | y | z |  |  |
| L posterior cingulate cortex | 31 | -6 | -44 | 38 | 7.26 | 0.001 |
| L inferior parietal lobule | 39 | -46 | -72 | 36 | 7.18 | 0.001 |
| R inferior parietal lobule | 39 | 56 | -64 | 20 | 6.02 | 0.016 |
| R ventral mPFC | 10 | 0 | 64 | -12 | 5.98 | 0.017 |
| L inferior temporal lobe | 21 | -52 | -16 | -18 | 6.59 | 0.004 |
| R inferior temporal lobe | 22 | 58 | -6 | -12 | 5.32 | 0.078 |
| L parahippocampal cortex | 36 | -24 | -20 | -20 | 8.18 | 0.000 |
| R parahippocampal cortex | 36 | 24 | -20 | -22 | 8.45 | 0.000 |

Whole brain random-effects analysis; $p < .05$ , FWE corrected, cluster size $\geq 10$ voxels.

### Resting state fMRI

Figure 3 shows a DMN activation at group level by FC analysis with rs-fMRI data. The peak coordinates of the PCC, the biliteral IPL, the vmPFC, the biliteral ITL and the biliteral PHC were used as the centers of the ROI spheres (radius 10 mm).

**Figure 3.**
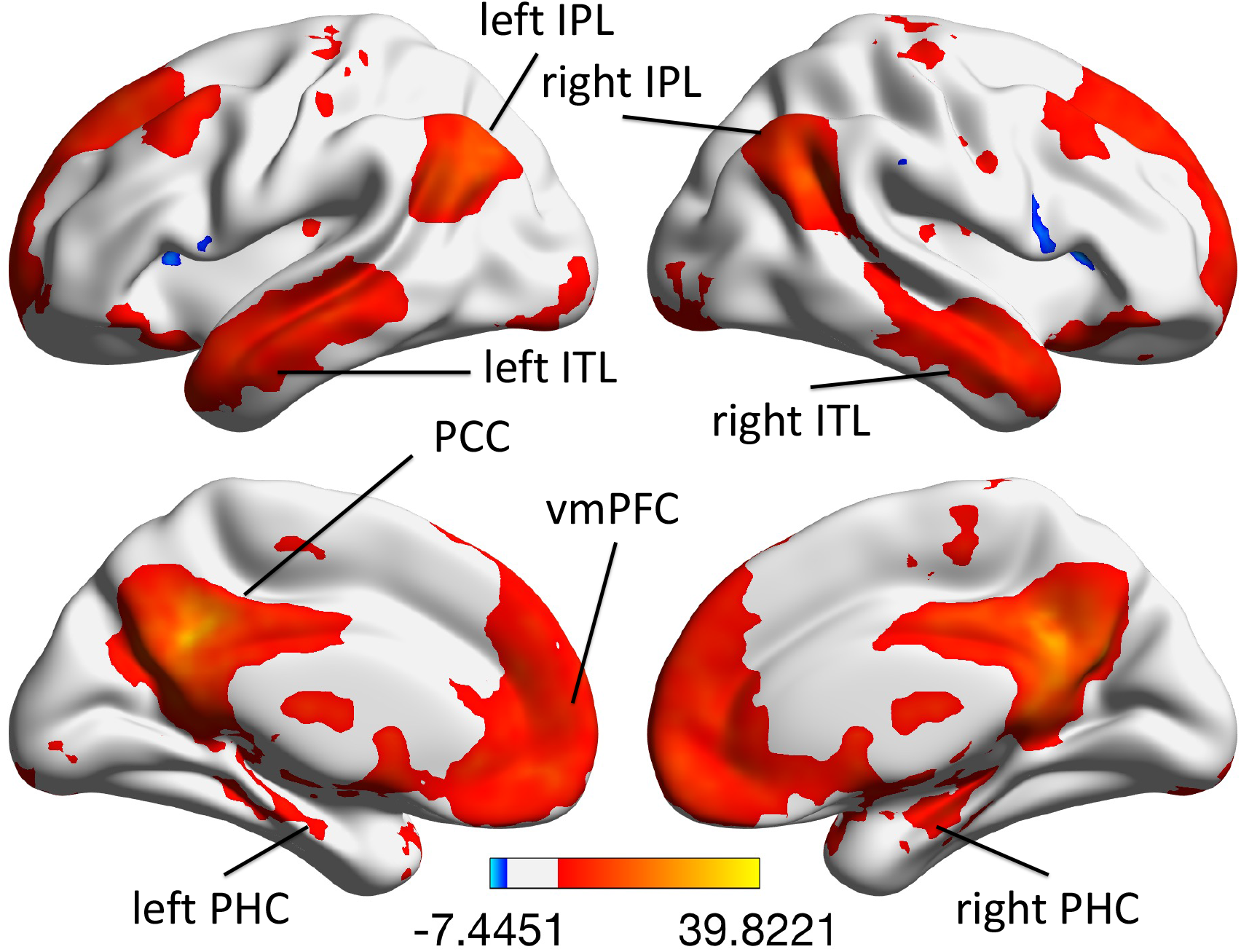
The DMN mapped by FC analysis with rs-fMRI data (*p* < .05, FWE corrected, cluster size ⩾ 10 voxels).

In Experiment 1 & 2, we conducted a same group GLM analysis contrasting the topological discrimination (based on the presence/absence of “hole”) and the local geometrical discrimination (based on orientation of angles/arrows, a kind of Euclidean property; parallelism, a kind of affine property; and collinearity, a kind of projective property) to identify brain regions involved in the processing of global topological properties. The results of the two experiments consistently revealed that the brain responses to the topological discrimination tasks were significant “above” that to the local discrimination tasks in a set of brain regions, which we named the Topological Discrimination Network (TDN). Further analysis demonstrated that the TDN not only has identical spatial distribution, but also essentially the same hemodynamic characteristics of functional signal responses with the DMN. Unsurprisingly, the discrimination of local geometrical properties exhibited a significant TID comparing with the fixation/resting period.

In Experiments 1 and 2, a total 8 sets of stimuli were adopted to reveal the brain response to topological discrimination in contrast with that to the local discrimination, 4 sets represented topological property and the rest represented different kinds of local geometrical properties. The consistent results make it very unlikely that the TDN resulted from responses to some specific stimuli or figures rather than reflecting the processing of topological properties.

## The Topological Discrimination Network Measured at 7T

### EXPERIMENT 3

#### METHODS

##### Subjects

7 males and 18 females, between 19 to 35 years old, participated in Experiment 3 as paid volunteers.

#### Stimuli and Procedure

The two sets of stimuli used in Experiment 1 (Figure 1A, stimulus ***a*** and ***b***) were tested at 7T MRI canner. Participants completed four 524-second blocked-design fMRI runs. Each run consisted of 16 blocks (half for each set of stimuli). Each block contained eight 2.5-second trials (1-second stimulus display and 1.5-second fixation). The tasks and the requirements for participants were all same as in the previous experiments at 3T scanner.

#### MR image acquisition and analysis

Imaging was performed at a Siemens 7T MEGNETON scanner (Erlangen, Germany) with a homemade 8-channel TX/RX head coil. A high-resolution EPI sequence was used for functional imaging [TR = 2,000 ms; TE = 23 ms; flip angle = 90°; parallel imaging with SENSE = 3; 40 descending axial slices; slice thickness = 2.0 mm; matrix size = 96 × 96; field of view (FOV) = 192 mm × 192 mm]. High-resolution T1-weighted 3D-MPRAGE images were acquired (TR/TE/TI = 2,200 ms/3.33 ms/1,050 ms; flip angle = 7°; 192 sagittal slices; field of view = 256 mm × 256 mm; resolution = 1.0 mm × 1.0 mm × 1.0 mm) for registration of functional images and localization of brain activation.

The same spatial pre-process (except unwarping) and 2-level statistics with the 3T experiments were conducted.

## RESULTS

Comparison of brain responses induced by the topological and local discrimination revealed relatively stronger responses to topological discrimination in the PCC, the vmPFC, the bilateral IPL, the bilateral anterior ITL and the bilateral PHC, which was consistent with the TDN observed in 3T experiments (Figure 4A). The time course of the BOLD responses to the topological discrimination were not significantly lower than that to the fixation condition in the bilateral ITL and the PHC (*p* > .05); the PCC, the vmPFC and the bilateral IPL showed a minor lower difference (*p* < .01, *p* = .493, *p* < .05, *p* < .05); the discrimination of local geometrical stimulus generated significant TID (*p* < .001) in all the regions compared with the resting (Figure 4B). All those activations passed the FWE correction (Table 3).

**Table 3.** The active regions of the topological vs. local geometrical discrimination

| Regions | BA | MNI coordinates | | | $t$ | $p$ |
| --- | --- | --- | --- | --- | --- | --- |
|  |  | x | y | z |  |  |
| L posterior cingulate cortex | 23 | 3 | -48 | 30 | 9.27 | 0.000 |
| L inferior parietal lobule | 39 | -50 | -69 | 36 | 10.75 | 0.000 |
| R inferior parietal lobule | 39 | 56 | -66 | 31 | 11.41 | 0.000 |
| R ventral mPFC | 32 | -8 | 45 | 1 | 9.88 | 0.000 |
| L inferior temporal lobe | 20 | -46 | -5 | -29 | 7.19 | 0.012 |
| R inferior temporal lobe | 38 | 44 | 10 | -33 | 7.61 | 0.006 |
| L parahippocampal cortex | 36 | -15 | -7 | -23 | 8.50 | 0.001 |
| R parahippocampal cortex | 36 | 20 | -10 | -23 | 8.09 | 0.002 |

Whole brain random-effects analysis; $p < .05$ , FWE corrected, cluster size $\geq 10$ voxels.

**Figure 4.**
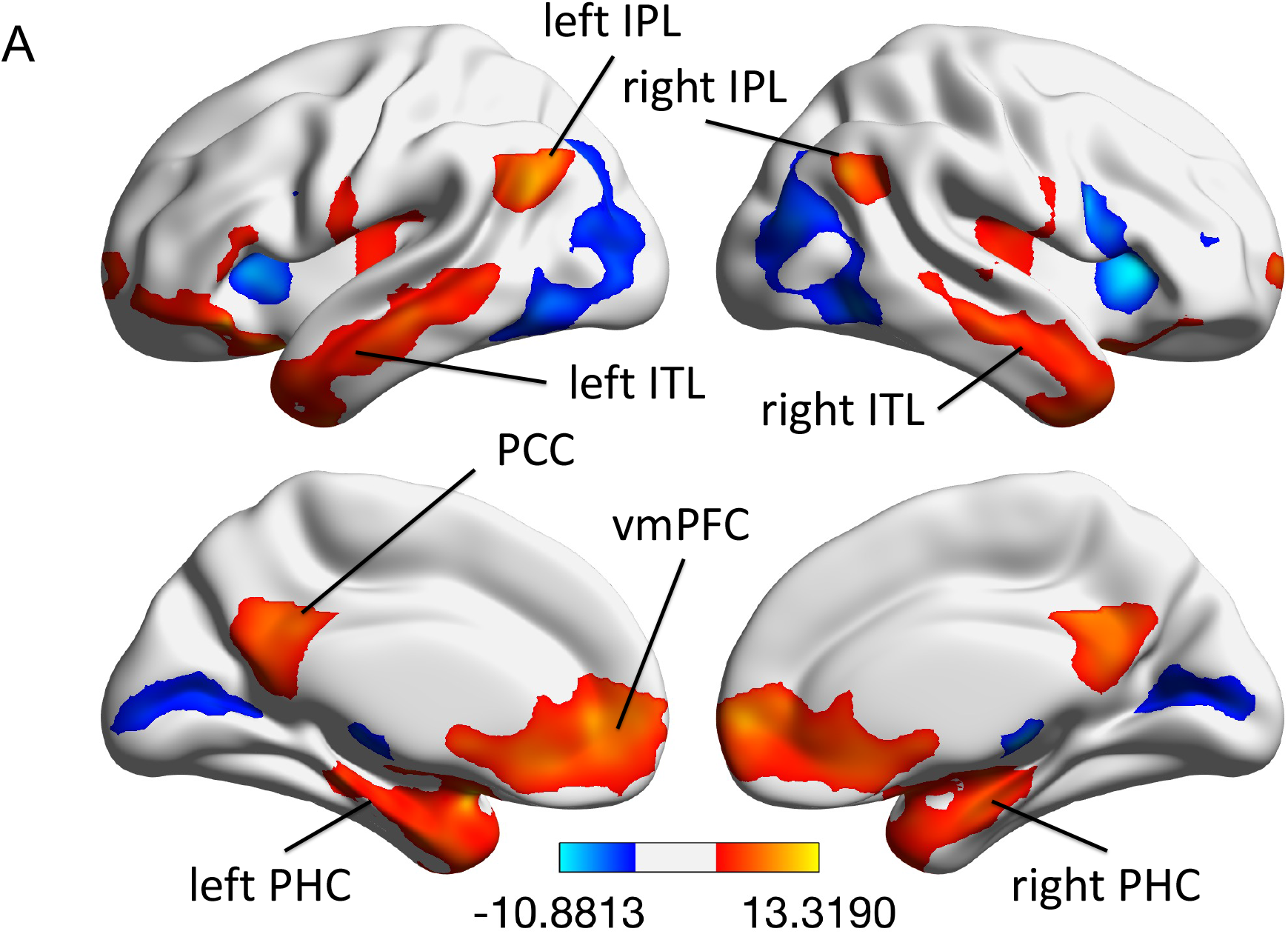
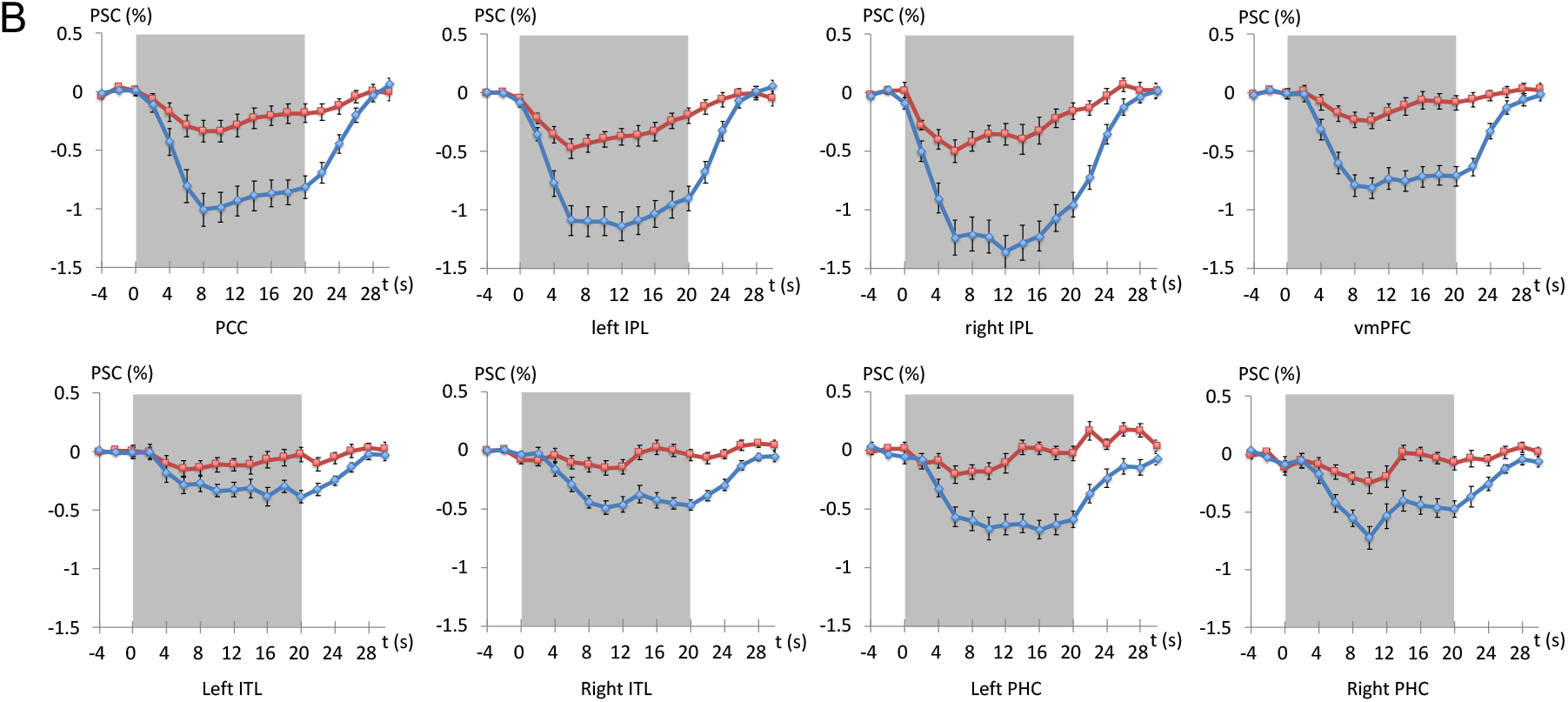
The results of Experiment 3. (A) comparison of fMRI activations between the topological and local discrimination tasks. (B) The averaged PSCs responding to the topological (red) and local (blue) tasks in the eight main ROIs. The gray box indicated the temporal duration of task blocks.

### EXPERIMENT 4

In addition to the “triangle-arrow pair” figure used in Experiment 1, stimulus ***d*** (Figure 2A) used in Experiment 2 is also a figure illustrating topological discrimination based on the difference in holes. In this experiment, we tested the stimulus and extended the functional imaging over whole brain with longer TR to verify the TDN at 7T scanner.

#### METHODS

##### Subjects

7 males and 19 females, between 20 to 25 years old, participated in Experiment 4 as paid volunteers.

#### Stimuli and Procedure

The two sets of stimuli used in Experiment 2 (Figure 2A, stimulus ***a*** and ***d***) were tested in this experiment. Participants went through five 261-second fMRI runs which consisted of six task blocks (half for each stimulus) and interleaved with 15-second fixation periods. Each task block contained eight 3.2-second trials. The stimulus display was presented for 1 second at the beginning of each trial. The tasks and requirements for participants were the same as in experiments 1∼3.

#### MR image acquisition and analysis

The same parameters except a longer TR (3,000 ms) and more slices (60 axial slices) than that used in Experiment 3 were adopted to achieve whole-brain EPI imaging. The same analysis as in Experiment 3 was conducted.

## RESULTS

### Voxel-wise analysis

The TDN was revealed by the comparison between topological and local orientation discriminations (Figure 5A). The time courses of the BOLD responses in the ROIs of the TDN showed highly consistent result with Experiment 3: the responses to topological discriminations were not significantly lower (in the bilateral ITL and the PHC, *p* > .05) or weakly lower (in the PCC, the bilateral IPL and the vmPFC, *p* < .05, *p* < .05, *p* < .05, *p* = .459) compared with the fixation. All activations passed FWE correction (Table 4).

**Table 4.** The active regions of the topological vs. local discrimination

| Regions | BA | MNI coordinates |  |  | *t* | *p* |
| --- | --- | --- | --- | --- | --- | --- |
|  |  | x | y | z |  |  |
| L posterior cingulate cortex | 23 | 8 | -48 | 33 | 12.30 | 0.000 |
| L inferior parietal lobule | 39 | -52 | -67 | 28 | 11.40 | 0.000 |
| R inferior parietal lobule | 39 | 54 | -61 | 34 | 11.41 | 0.000 |
| R ventral mPFC | 10 | -14 | 62 | 9 | 15.70 | 0.000 |
| L inferior temporal lobe | 38 | -46 | 9 | -38 | 10.60 | 0.000 |
| R inferior temporal lobe | 38 | 56 | 8 | -27 | 8.16 | 0.002 |
| L parahippocampal cortex | 36 | -24 | -19 | -18 | 8.69 | 0.001 |
| R parahippocampal cortex | 36 | 21 | -6 | -20 | 10.86 | 0.000 |

Whole brain random-effects analysis; $p < .05$ , FWE corrected, cluster size $\geq 10$ voxels.

**Figure 5.**
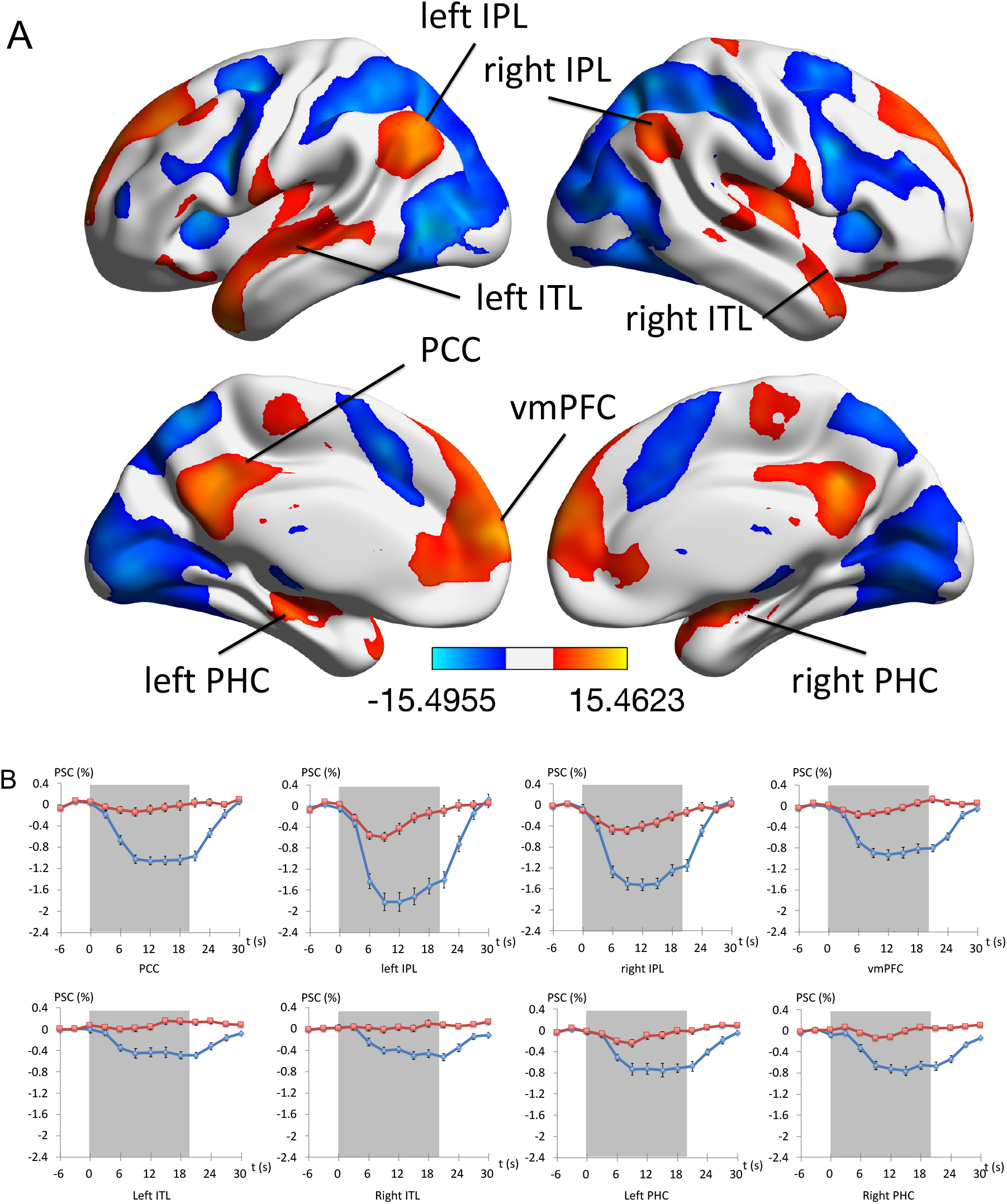
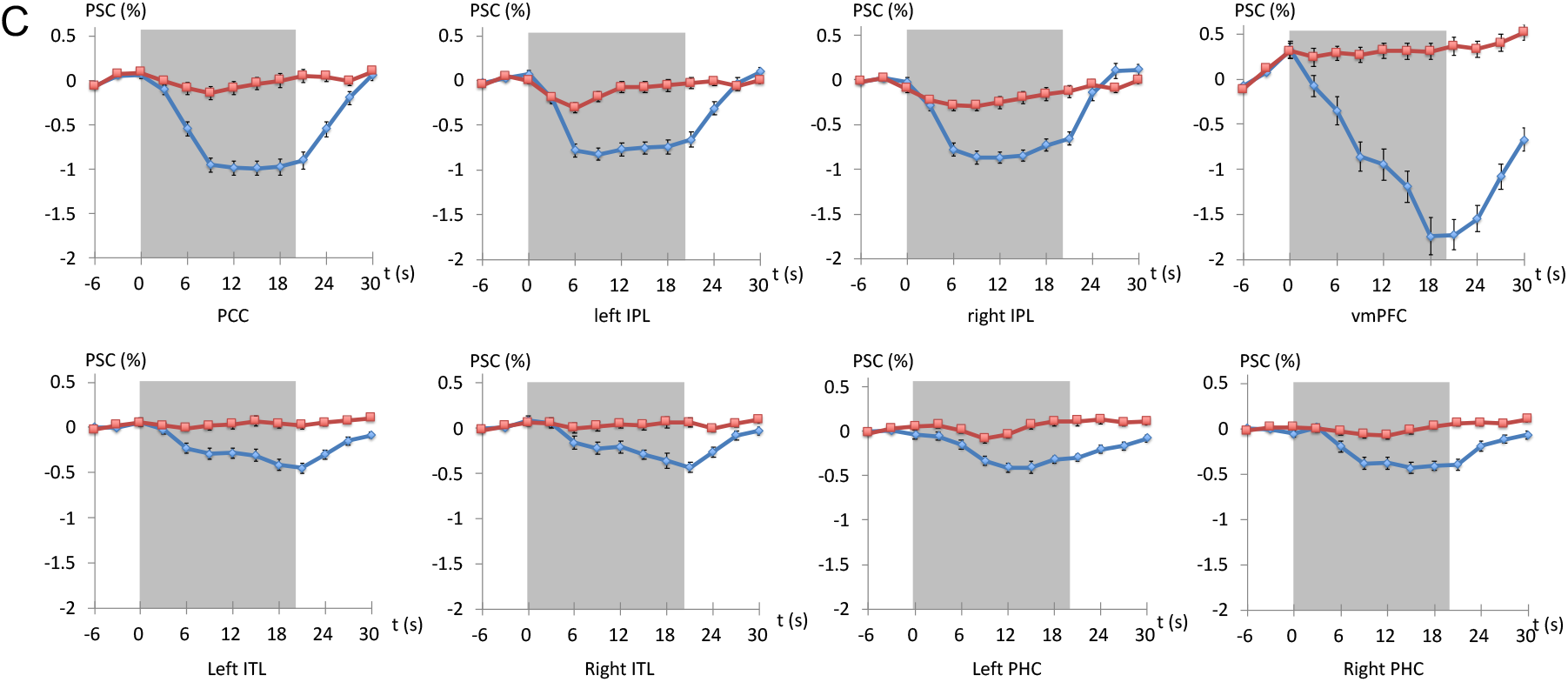
The results of Experiment 4. (A) fMRI activation patterns from the contrast of topological vs. local discrimination. (B) The average PSCs responding to topological (red) or local (blue) discrimination in eight major ROIs based on contrast in A. (C) The average PSCs responding to topological (red) or local (blue) discrimination in eight ROIs defined by independent rs-fMRI data. The gray box indicated the temporal duration of task blocks.

### ROI analysis

As shown in Figure 5C, in the ROIs defined by the resting-state fMRI (Figure 3), the PCC, the bilateral ITL and PHC, the BOLD responses to the topological discrimination task were not significantly different with that to the fixation (*p* > .05); in the vmPFC, the BOLD response to the topological discrimination task was higher than to the fixation (*p* < .001); in the bilateral IPL, or weakly lower than the fixation (*p* = .052; *p* < .005). The local orientation discrimination evoked a significant TID in all the ROIs compared with the fixation (*p* < .001).

fMRI at 7T provides higher signal to noise ratio (SNR) and contrast to noise ratio (CNR) than 3T, more importantly, signals obtained at 7T are more specific to neural responses because they are less influence by large draining veins. Consequently, based on the 7T data, the active regions revealed by the contrast of the topological and local discrimination task all passed the FWE correction (having equal sample size of the functional measurements at 3T). The functional responses to the topological discrimination did not any significant difference comparing with the resting/fixation condition in all ROIs of the TDN or the DMN, while the local discrimination induced a significant TID. The convergent results of 3T and 7T experiments further support the consistency of the TDN with the DMN.

## The TDN defined by Cerebral Blood Flow Changes

### EXPERIMENT 5

#### METHODS

##### Subjects

10 males and 8 females, between 21 to 28 years old, participated in Experiment 5 as paid volunteers.

#### Stimuli and Procedure

The stimuli were the same with Experiment 2. Participants completed 6 perfusion fMRI runs. Each run began with an 8-second fixation, followed by six 72-second task blocks (one block for each set of stimuli) and interleaved with 72-second fixation. Each block contained twenty-four 3-second trials. The stimulus display was presented for 2.5 seconds at the beginning of each trial. The task and requirements for participants were the same as in the BOLD-based experiments.

#### MR image acquisition and analysis

Participants were scanned by a Siemens 3T Trio scanner (Erlangen, Germany) with a single-channel receive-only head coil. ASL perfusion MRI was performed with continuous arterial spin labeling (CASL) sequence (single-shot gradient-echo echo-planar imaging, TR/TE = 3000/17 ms, flip angle = 90°; 12 ascending axial slices, slice thickness = 8.0 mm; matrix = 64 × 64; FOV = 220 mm × 220 mm; labeling duration = 1650 ms, post labeling delay = 1200 ms). The same high-resolution T1-weighted image with previous experiments at 3T was acquired.

The first two EPI images of each run were discarded to allow for magnetization equilibration. The rest EPI images were realigned, co-registered to the structure image and smoothed using a 10 mm FWHM isotropic Gaussian kernel with SPM8. Cerebral blood flow (CBF) images were reconstructed and averaged to mean images by each stimulus. Then the mean CBF images were normalized to MNI space. Pair *t* test was conducted to discover the difference of regional CBF (rCBF) between different tasks. The global-mean scaling was done for eliminating the baseline differences of CBF across subjects, by using the proportional normalization method in the GLM model.

## RESULTS

### Voxel-wise analysis

The results demonstrated a consistent patterns of activation with the BOLD-based fMRI experiments 1∼4: The topological discrimination caused significantly higher rCBF than local one in the PCC, the vmPFC, the bilateral IPL and ITL (Figure 6A, Table 5), which were all part of the TDN or the DMN. No significant rCBF deficit was observed comparing the topological discrimination task with the fixation period in those regions (*p* > .05), except the bilateral IPL (*p* < .05). As a contrast, the local discrimination task induced significant CBF deficit in those regions compared with the fixation.

**Table 5.** The active regions of the topological vs. local discrimination

| Regions | BA | MNI coordinates | | | $t$ | $p_{un}$ |
| --- | --- | --- | --- | --- | --- | --- |
|  |  | x | y | z |  |  |
| L posterior cingulate cortex | 23 | 12 | -46 | 34 | 9.27 | 0.000 |
| L inferior parietal lobule | 39 | -44 | -74 | 40 | 10.75 | 0.000 |
| R inferior parietal lobule | 39 | 56 | -58 | 42 | 11.41 | 0.000 |
| R ventral mPFC | 11 | 4 | 62 | -18 | 9.88 | 0.000 |
| L inferior temporal lobe | 21 | -62 | -12 | -24 | 7.19 | 0.012 |
| R inferior temporal lobe | 20 | 58 | -14 | -30 | 7.61 | 0.006 |

Whole brain random-effects analysis; $p < .001$ , uncorrected, cluster size $\geq 40$ voxels.

**Figure 6.**
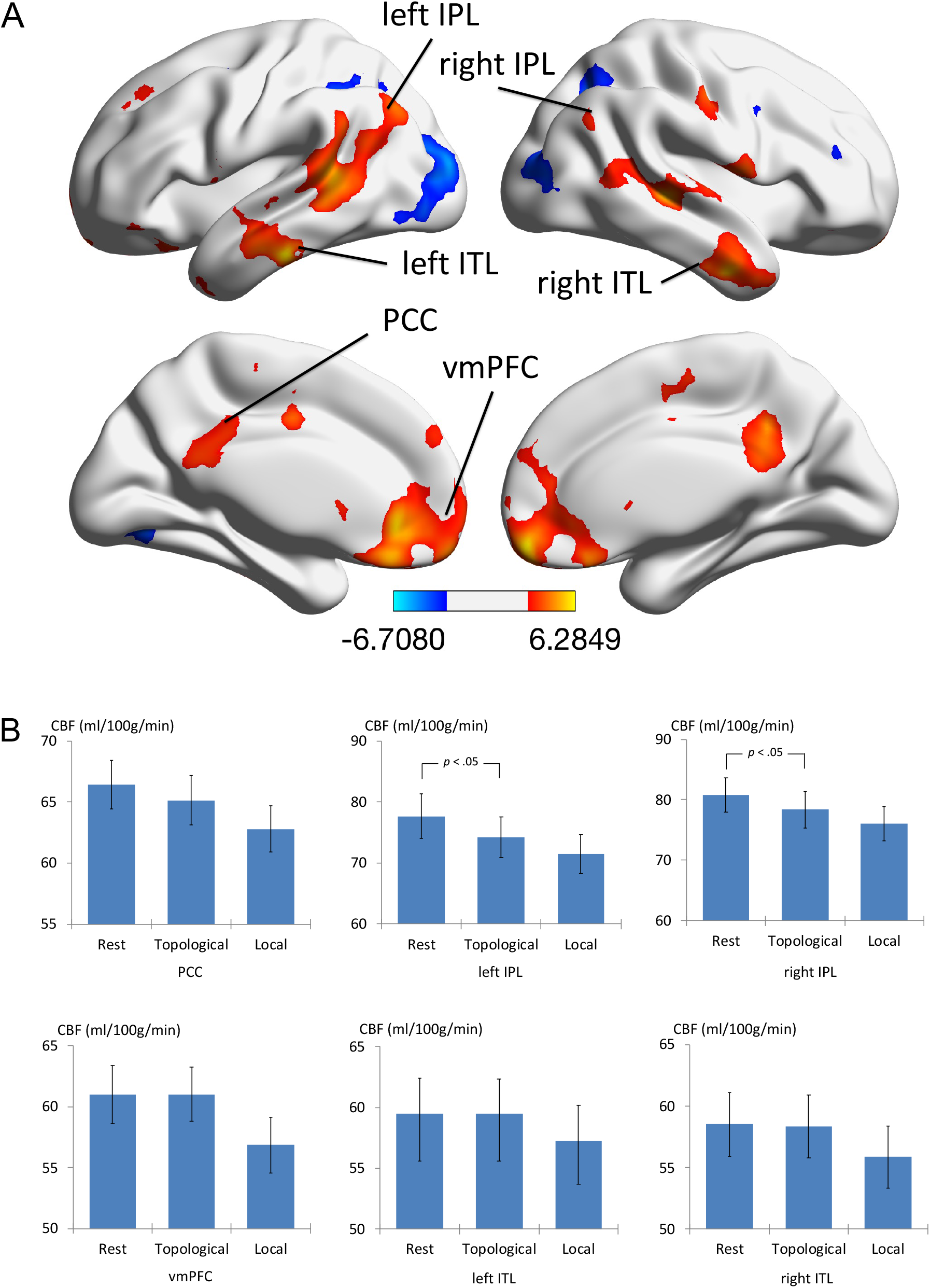
Results of Experiment 5. (A) rCBF activation patterns contrasting the topological vs. local discrimination tasks (*p* < .001, uncorrected, cluster size ≥ 40). (B) The average rCBF values of six main active regions in A.

### EXPERIMENT 6

In this experiment, a new pair of “S-like” and “ring” figure was adopted to represent the topological difference in holes. And a new pseudo-continuous arterial spin labeling (pCASL) imaging technique(Alsop et al., 2015) was used in CBF measurement to improve the SNR.

#### METHODS

##### Subjects

8 males and 13 females, between 20 to 28 years old, participated in Experiment 6 as paid volunteers.

#### Stimuli and Procedure

A new set of stimuli were used in this test. Stimulus ***a*** (Figure 7Aa) served as a discrimination task based on mirror-asymmetry of S-like figure, also a kind of Euclidean property. Stimulus ***b*** (Figure 7Ab) illustrated topological discrimination based on the difference in holes by using an S-like figure vs. a ring. This S-like figure and the ring were designed to control for luminous flux, spatial frequency components, and other possible confounds of local features(Chen, 1990; Chen et al., 2003). The S-like figure was scaled to approximate the area of the ring, and its shape was purposely made irregular in order to eliminate the possible effects of subjective contours or other organizational factors (such as parallelism, or similarity of length). As a consequence, the S-like figure and the ring differ in holes, but are quite similar in local features, such as luminous flux, spatial frequency components, perimeter length, and averaged edge crossings. The small figure in each quadrant and the whole stimulus were displayed in size of 5.0° × 5.0° and 10.8° × 10.8°, respectively.

**Figure 7.**
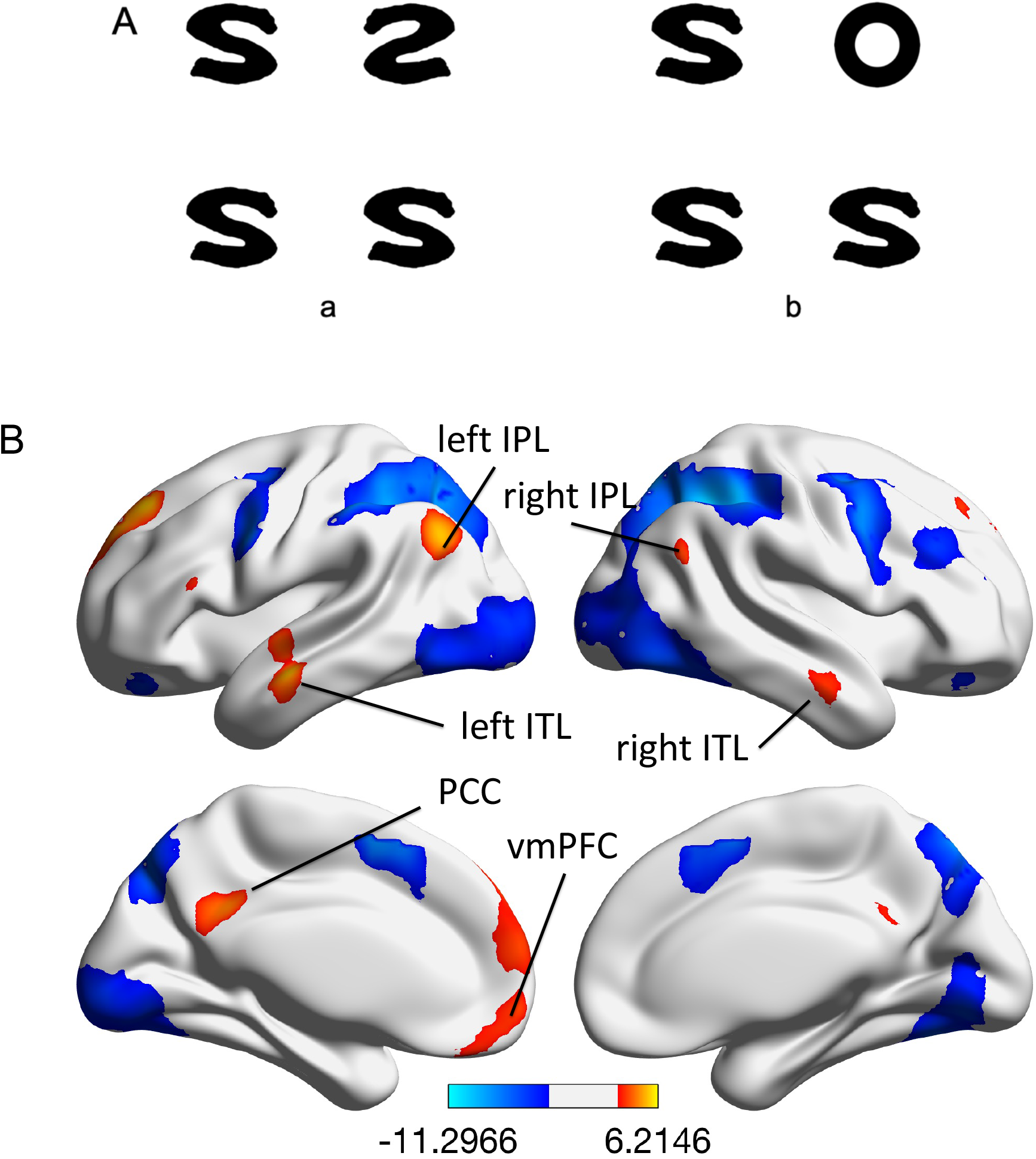
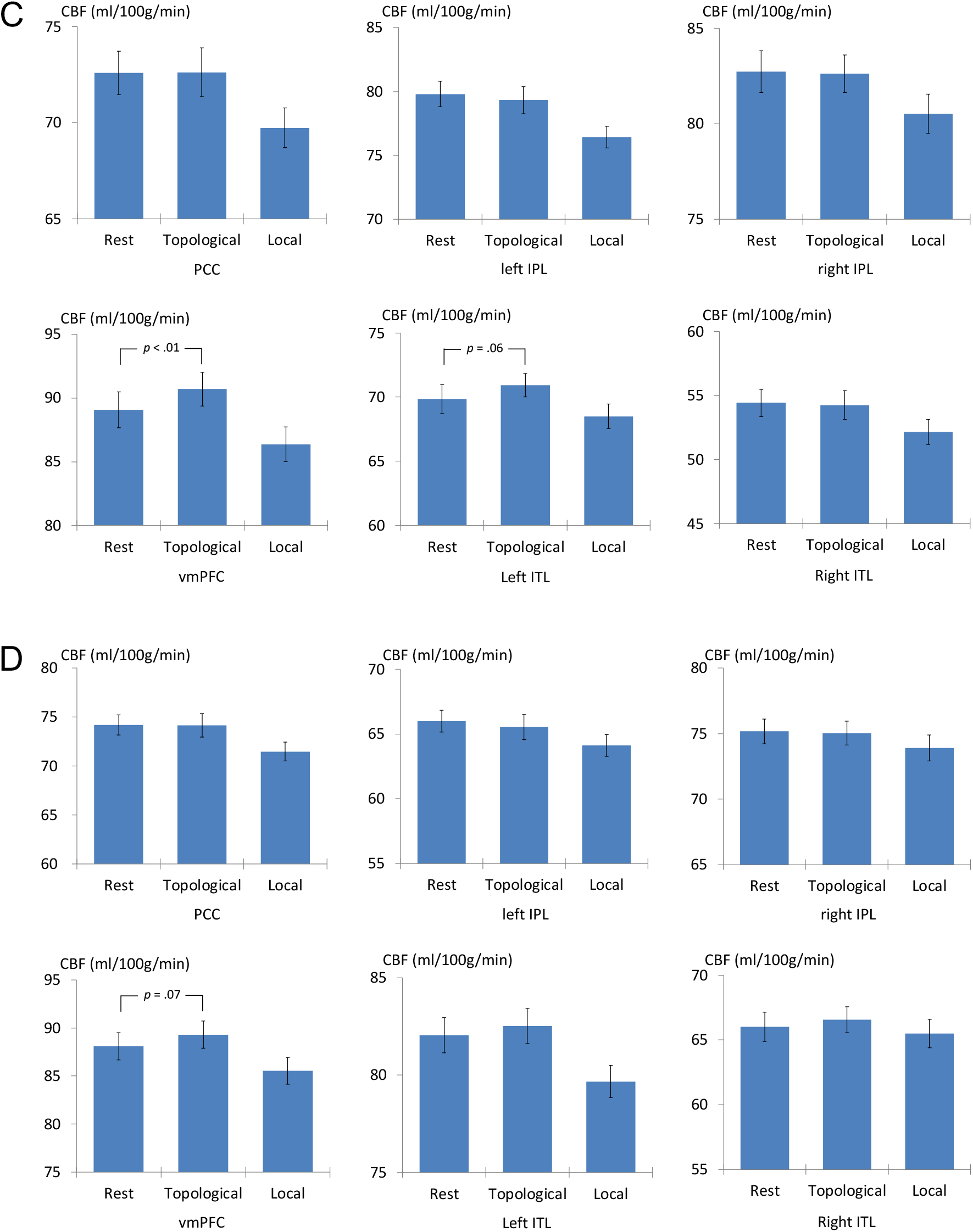
The stimuli and results of Experiment 6. (A) The two sets of stimuli used in this experiment; (B) rCBF activations of the topological vs. local discrimination (*p* < .001, uncorrected, cluster size ≥ 40). (C) The average rCBF values of six main active regions in B. (D) The average rCBF values of the six ROIs defined by resting-state fMRI data.

Participants completed 4 runs. Each run consisted of eight task blocks and each block contained eight 2.5-second trials. The stimulus display was presented for 1 second at the beginning of the trial. The task, the procedure and the requirements to the subjects were the same with previous experiments 1∼5.

#### MR image acquisition and analysis

The subjects were scanned by a Siemens 3T Trio Tim scanner (Erlangen, Germany) with a 12-channel receive-only head coil. The CBF images were acquired with a pCASL sequence (TR/TE = 4000/17.96 ms, flip angle = 90°; parallel imaging with SENSE = 3; 24 ascending axial slices, slice thickness = 5.0 mm; matrix = 64 × 64; FOV = 192 mm × 192 mm; labeling duration = 1800 ms, post labeling delay = 1800 ms). The structure image of the whole brain was acquired with the same sequence with the 3T experiments 1∼2. The same analysis with Experiment 5 was conducted.

## RESULTS

### Voxel-wise analysis

The result was consistent with Experiment 5: six active regions (Table 6) revealed by topological discrimination vs. mirror asymmetry, all fell in the TDN (Figure 7B). There was not significant rCBF deficit of the topological discrimination task compared with the fixation in any of the regions (*p* > .05). A significant increase of rCBF was observed in the vmPFC and the left ITL (*p* < .01, *p* = .06) when topological discrimination task was performed.

**Table 6.** The active regions of the topological vs. local discrimination

| Regions | BA | MNI coordinates |  |  | *t* | *p<sub>un</sub>* |
| --- | --- | --- | --- | --- | --- | --- |
|  |  | x | y | z |  |  |
| L posterior cingulate cortex | 23 | -6 | -48 | 32 | 10.41 | 0.000 |
| L inferior parietal lobule | 39 | -48 | -68 | 36 | 11.48 | 0.000 |
| R inferior parietal lobule | 39 | 56 | -64 | 28 | 8.57 | 0.001 |
| R ventral mPFC | 10 | -4 | 60 | -6 | 8.24 | 0.002 |
| L inferior temporal lobe | 21 | -58 | -12 | -14 | 6.59 | 0.004 |
| R inferior temporal lobe | 21 | 58 | -8 | -24 | 13.59 | 0.000 |

Whole brain random-effects analysis; $p < .001$ , uncorrected, cluster size $\geq 40$ voxels.

### ROI analysis

Compared with the fixation, no significant deficit of rCBF response to the topological discrimination task was observed in all resting-state fMRI defined ROIs (*p* > .05). In contrast, the local discrimination task induced significant rCBF deficits compared with fixation. The current CBF results are consistent with the results of the BOLD-based fMRI experiments 1∼4.

CBF-based fMRI has more acute responses to the specific cognitive tasks because of its more directly coupling with the neural activity than classic BOLD fMRI (Detre, Rao, Wang, Chen, & Wang, 2012; Jiongjiong Wang et al., 2003), albeit with limitations on temporal and spatial resolution. Based on it, the rCBF measured during the topological discrimination task showed no difference with the fixation, providing robust biological evidence for the consistency between the TDN and the DMN.

## General Discussion

In this study, we found that a task-evoked network, the TDN, possesses the characteristics of the DMN: (1) the TDN’s hubs and the DMN’s hubs are well matched, (2) all non-topological tasks (discrimination of local geometric properties) led to deactivations at these hubs, and (3) the time courses of BOLD as well as the regional CBF signals in the TDN’s hubs are largely the same as in the DMN’s. In other words, the task-evoked TDN is consistent with the resting DMN. Such an observation could help us to understand the key question of “what is the function of the resting DMN”.

### Little or no deactivation in the TDN/DMN hubs from the topological discrimination tasks

The default mode hypothesis is based on the repeated observation that certain brain areas show TID across almost all of cognitive tasks. These decreases suggest the existence of an organized, baseline default mode of brain function that is suspended during specific goal-directed behaviors. In the current study, we tested this hypothesis by comparing the BOLD or the CBF activations in response to the topological and local geometric discrimination task. The analyses revealed three important findings. First, the areas defined by topological vs. local discrimination tasks, named TDN, consistently matched that of the DMN. Second, the TDN showed a resting-like responses to the topological discrimination task with minimal level of activation or deactivation. Third, local geometric discrimination tasks induced significant and sustained deactivation in the TDN.

### Baseline but not resting

In cognitive neuroscience, the designation of a condition as the control or the baseline, so that it could be compared against by the condition of interest, has been a very important issue. Defining a baseline state in the human brain, arguably our most complex system, poses a particular challenge(Debra A Gusnard & Raichle, 2001; Raichle et al., 2001), although researchers have treated fixation (or eye-closed) as a natural baseline in functional brain imaging. However, treating “fixation” or “eyes-closed” as the baseline control raises serious issues because of the lack of real control for eyes-closed or visual fixation in the so-called “resting” state. In this study, we observed that the dynamic changes of the BOLD signals during topological discrimination task were essentially the same as during the fixation phase, while pronounced TID were caused by local geometric discrimination tasks. This is the first time that a discrimination task not evoking a TID was observed in the main hubs of DMN. Thus the topological discrimination task, instead of “resting”, could potential serve as a cognitive baseline for various perceptual/cognitive tasks, with the advantage of having definite operational variables and well-controlled paradigms.

### Consistency of TDN across field strengths and signal types

The definition and characterization of TDN in the current study were achieved at both 3T and 7T, with BOLD-based as well as CBF-based signals. Higher field strength at 7T helped to reduce the influence of venous signals on fMRI measurements and increase the SNR (Kalcher, Boubela, Huf, Našel, & Moser, 2015). Our two 7T experiments, while showed fundamentally the same pattern of TDN, did improve the accuracy of functional localization and the temporal stability of BOLD responses in the TDN.

As a powerful complementary measure to the BOLD fMRI, ASL MRI technique has played a dominated role in CBF-based fMRI field due to its safety, simplicity, cost efficiency, and excellent reproducibility over longer time periods, etc. CBF measures have been used in brain activation studies as a well-established correlate of brain function and are considered to be more directly coupled with the neural activity than BOLD. Because of its close coupling with neural activity and its tolerance to magnetic-field inhomogeneity effects, the stimulus-driven acute CBF responses could play an important role for investigating brain functions (Aguirre, Detre, Zarahn, & Alsop, 2002; Ching-Mei Feng, 2004; J. Wang et al., 2003) (J. Wang et al., 2004). It is reassuring that in experiments 5 and 6 of the current study, the CBF changes obtained from ASL perfusion fMRI technique also revealed the TDN, providing converging evidence for the consistency between the TDN and the DMN.

## Conclusion

Although the DMN has been identified reliably during rest and is considered to reflect the intrinsic functions of the brain, its basic function remains unclear. With multi-modal functional imaging at multiple field strengths, our findings that the TDN was spatially consistent with the DMN. We believe that the TDN/DMN might be involved in the processing of topological properties. An important implication of this finding is that the TDN would be conducted as a general well-controlled baseline in cognitive fMRI investigations, instead of relying on “fixation” or “eye-closed resting” as the baseline.

## Acknowledgments

The authors would like to thank Yan Zhuo, Jing Luo and Zentao Zuo for their advices on data acquisition, and Ke Zhou and Hengyi Rao for their help in data analysis, interpretation, and manuscript writing. The project was supported by the Ministry of Science and Technology of China grant (2022ZD0211901, 2019YFA0707103, 2020AAA0105601), the National Nature Science Foundation of China grant (31730039,U21A20388), and the Chinese Academy of Sciences grants (ZDBS-LY-SM028).

